## Supplementary information for "Subcortical Circuits Mediate Communication Between Primary Sensory Cortical Areas"

Lohse et al.

### Supplementary information

Supplementary Figures 1-14

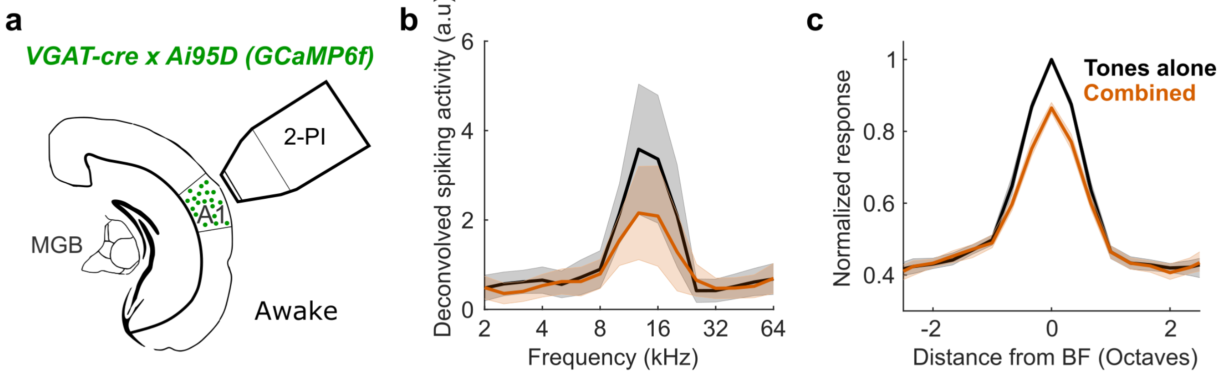


**Supplementary Fig. 1 – Related to Figures 1 and 4. Inhibitory interneurons in A1 are suppressed by whisker stimulation. a** Schematic of experimental setup in which 2-photon calcium imaging of VGAT+ cells in A1 was performed while concurrently presenting tones and stimulating the whiskers**. b** Frequency response profile (at 60 dB SPL) of an example VGAT+ cell in A1 with (orange) or without (black) concurrent whisker deflection. **c** Median frequency response profile across all frequency-tuned VGAT+ cells in A1 with (orange) or without (black) concurrent whisker deflection ($n$ = 514 cells, 3 mice). Shading illustrates the 95% confidence intervals around the mean (**b**) or nonparametric 95% confidence intervals around the median firing rate (**c**).

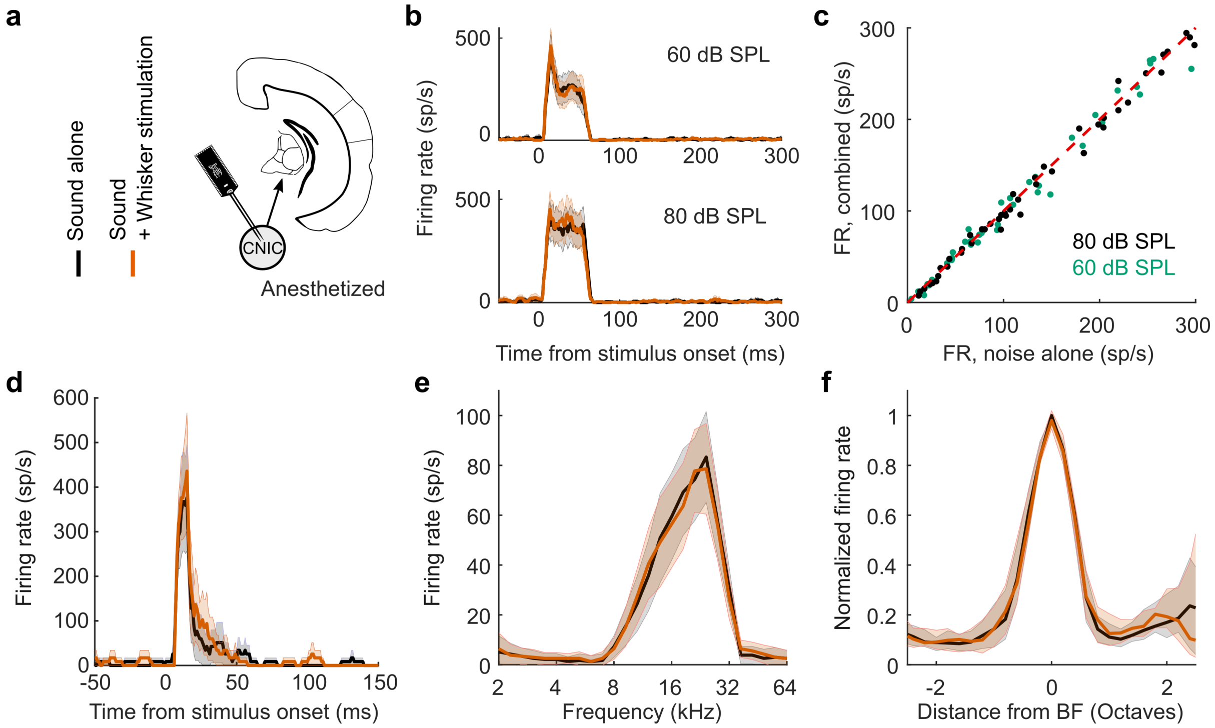


**Supplementary Figure 2 – Related to Figure 2. Somatosensory modulation of auditory processing is absent from the central nucleus of inferior colliculus (CNIC) in anesthetized mice. a** Schematic of recording setup. **b** Example CNIC unit responses to broadband noise (at either 60 or 80 dB SPL) with (orange) or without (black) concurrent whisker stimulation. **c** Summary of firing rate (FR) responses of CNIC units to broadband noise alone or combined with concurrent whisker stimulation (Wilcoxon signed rank tests, P > 0.05, $n_{60dB}$ = 45, $n_{80dB}$ = 48, 2 mice). **d** Example CNIC unit response to a tone presented at its BF at 80 dB SPL with (orange) or without (black) concurrent whisker deflection. **e** Frequency response profile for the same CNIC unit with (orange) or without (black) concurrent whisker deflection. **f** Median frequency response profiles across all frequency tuned CNIC units with (orange) or without (black) concurrent whisker deflection (*n* = 58, 2 mice). Shading illustrates the s.e.m. (**b**,**d**), the 95% confidence intervals around the mean (**e**) or nonparametric 95% confidence intervals around the median firing rate (**f**).

To assess whether the somatosensory effects on the auditory thalamocortical system are relayed from an earlier structure, prior to thalamus, in the auditory pathway, we recorded extracellular unit activity from the CNIC, which delivers the bulk of the auditory input to MGBv. These recordings showed no effect of whisker stimulation on responses to tones or broadband noise in the CNIC. Similarly, whisker stimulation alone did not affect the baseline firing rate across the population of noise-responsive units in IC (P > 0.05, Wilcoxon signed-rank test, *n* = 48). In accordance with this, the proportion of CNIC units in which a significant (P < 0.05, *t*-test) change in firing was observed during whisker stimulation was no different from what would be expected by chance (P > 0.05, binomial test). However, it is worth noting that the few units showing a significant increase in firing rate (3/48) were histologically localized to the lateral part of IC, which receives direct projections from S1 (see Fig. 7).

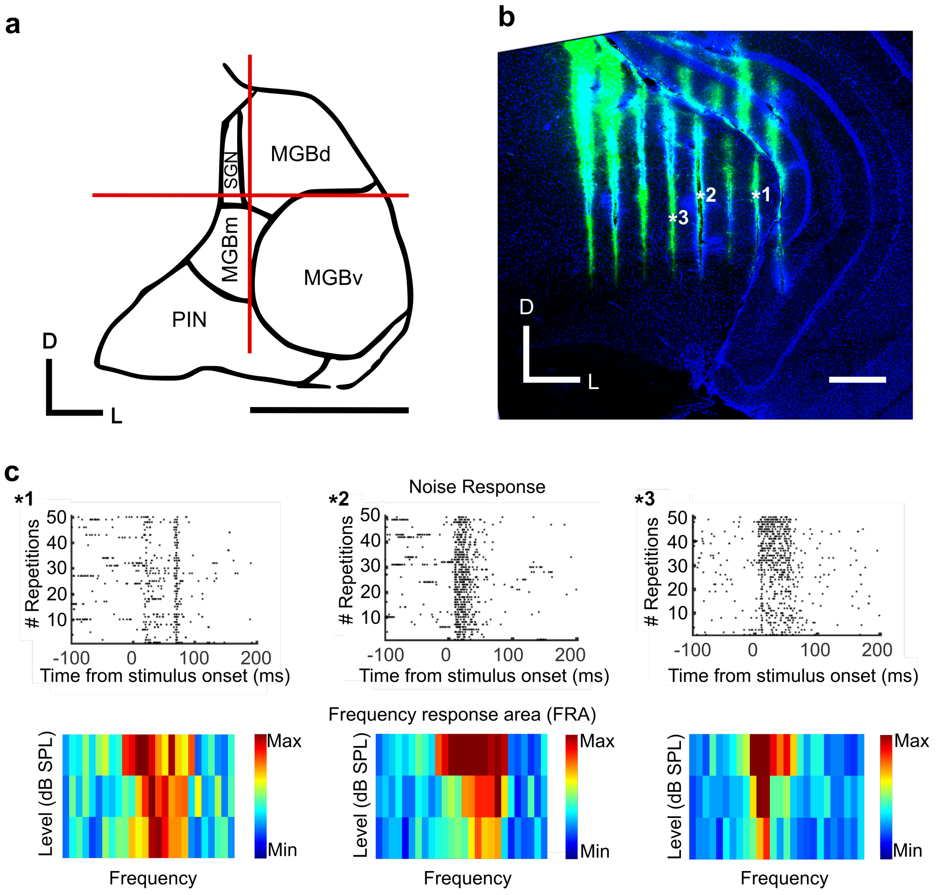


**Supplementary Figure 3 – Related to Figure 2. Multiunit activity and histological identification of recording sites in auditory thalamus of anesthetized mice. a** Schematic of auditory thalamus (based on Lu et al., 2009). Red lines indicate the grid used to allocate recording sites to auditory thalamus subdivisions. Scale bar, 500 µm. D, dorsal; L, lateral. **b** Confocal image of a coronal section at the level of the MGB with DiI traces (displayed as green) left by an 8 x 8 configuration linear Neuronexus probe. Scale bar, 400 µm. **c** Example raster plots to 80 dB SPL broadband noise stimulation (top) and frequency response areas (bottom, 20 dB attenuation steps, 4-64 kHz) from multiunit activity recorded at the locations marked in **b**.

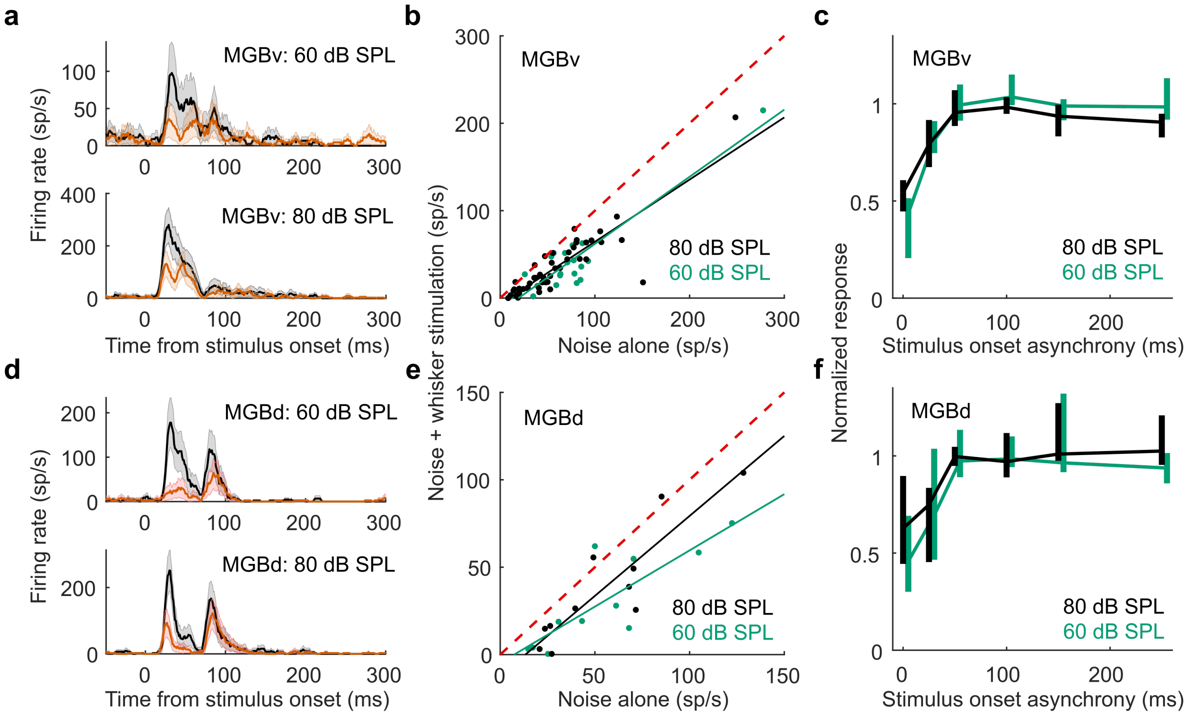


**Supplementary Figure 4 – Related to Figure 2. Temporal dependence of somatosensory suppression of responses to broadband noise in the MGBv and MGBd of anesthetized mice. a** Examples unit responses in MGBv to 60 dB SPL (*top*) and 80 dB SPL (*bottom*) broadband noise either with (orange) or without (black) concurrent whisker stimulation. Shading illustrates the s.e.m. around the means. **b** Summary of MGBv responses to 60 dB SPL or 80 dB SPL broadband noise with vs without concurrent whisker stimulation. **c** Median somatosensory suppression of auditory responses as a function of increasing stimulus onset asynchronies in MGBv, $n_{80dB\_SPL}$ = 47 (6 mice), $n_{60dB\_SPL}$ = 22 (5 mice). Error bars are 95% bootstrapped nonparametric confidence intervals around the median. **d-f** As in **a-c**, but for units recorded in MGBd, $n_{80dB\_SPL}$ = 11 (3 mice), $n_{60dB\_SPL}$ = 11 (3 mice). Median $change_{MGBv\_80dB}$ = -45.6%, $P_{MGBv\_80dB} < 0.001$; median $change_{MGBd\_80dB}$ = -37.3%, $P_{MGBd\_80dB} < 0.001$; median $change_{MGBv\_60dB}$ = -56.4%, $P_{MGBv\_60dB} < 0.001$; median $change_{MGBd\_60dB}$ = -54.3%, $P_{MGBd\_60dB} < 0.001$, Wilcoxon signed rank tests).

Somatosensory suppressive effects on auditory processing were still significant when the noise onset was delayed by 25 ms relative to the whisker deflection onset for both 60 dB SPL and 80 dB SPL broadband noise stimuli ($P < 0.001$, Wilcoxon signed rank tests), but not when the noise onset occurred 50 ms or more after the onset of the whisker stimulation. Because whisker stimulation lasted 50 ms, the suppressive effects on MGBv were present as long as the sound was presented during, but not outside, the whisker stimulation window. A similar effect of offsetting the noise stimulus was found for MGBd, although somatosensory suppressive effects on auditory responses were no longer significant when the 60 dB SPL stimulus was delayed by 25 ms, possibly due to the small sample size ($P = 0.067$, Wilcoxon signed rank test). Whisker stimulation did not affect baseline activity in noise-responsive units in either MGBv ($P > 0.05$, $n$ = 47, 6 mice, Wilcoxon signed rank test) or MGBd ($P > 0.05$, $n$ = 11, 3 mice, Wilcoxon signed rank test).

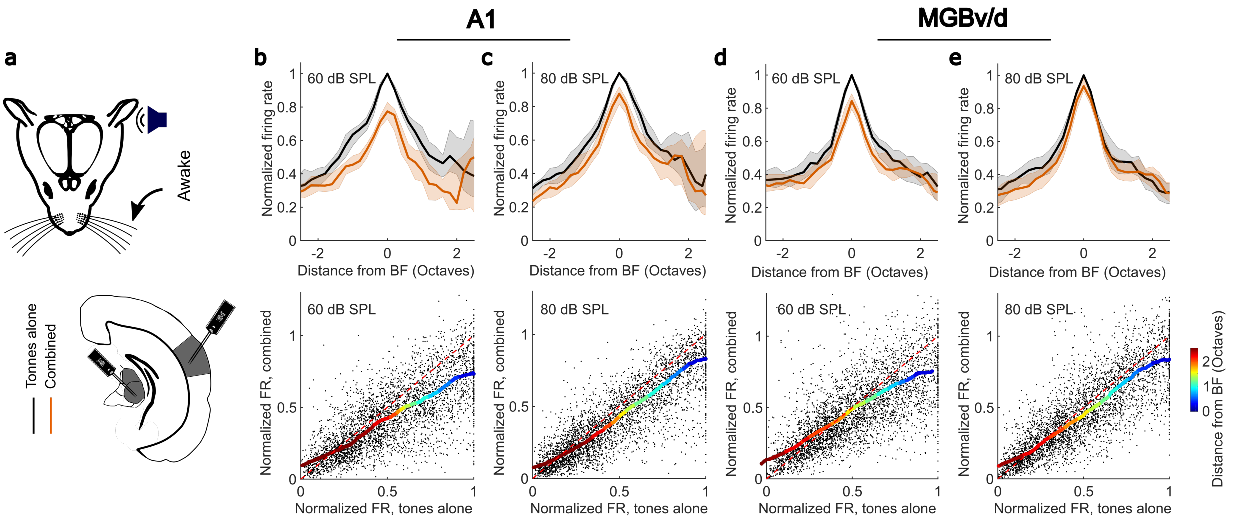


**Supplementary Fig. 5 – Related to Figure 2. Somatosensory suppression in A1 and MGBv/d of awake, passively listening mice. a** Schematic of recording setup. **b**,**c** *Top:* Median frequency response profiles for tones presented at 60 (**b**) or 80 dB SPL (**c**) across units recorded in A1 of awake mice (60 dB SPL change in BF response: P < 0.001, *n* = 140; 80 dB SPL change in BF response: P < 0.001, *n* = 140). *Bottom:* Relationship between normalized firing rate (FR) for all units (black dots) recorded in A1 in response to tones presented at 60 (**b**) or 80 dB SPL (**c**) across all frequencies presented either with ('combined') or without ('tones alone') whisker stimulation. Thick multi-colored lines show the running median of this relationship (window: 0.1 normalized firing rate), and the colors denote distance from BF. The diagonal dashed red line is the line of equality. A larger distance between the multi-colored line and the diagonal line at the blue end than at the red end indicates divisive scaling. **d**,**e** Same as panels **b**,**c** but for units recorded in MGBv or MGBd (60 dB SPL change in BF response: P < 0.001, *n* = 157; 80 dB SPL change in BF response: P = 0.01, *n* = 157). Shaded areas in **b-e** indicate the 95% nonparametric confidence intervals of the median. $n_{A1}$ = 140 (4 mice); $n_{MGBv/d}$ = 157 (5 mice). Note that the suppressive effects of whisker stimulation on auditory responses showed the same sound level dependence in MGBv/d and A1.

In awake mice, a small subset of noise-responsive units in A1 (13.9%, *n* = 115, which is a higher proportion than expected by chance, P = 0.004, binomial test) showed significant (P < 0.05, *t*-test) firing rate changes in response to whisker stimulation alone in both directions (10.4% decreasing firing rate, and 3.5% increasing firing rate). In anesthetized mice, we also found that a subset of units in A1 (25%, *n* = 80, P < 0.001, binomial test) showed significant firing rate changes in response to whisker stimulation alone, again in both directions (5% decreasing firing rate, and 20% increasing firing rate).

In MGBv/d of awake mice, a small but significant proportion (12.8%, *n* = 47, P = 0.03, binomial test) of units in MGBv showed a significant increase in firing in response to whisker stimulation alone. In anesthetized mice, we also found a subset of units in MGBv/d (8.8%, *n* = 58, P = 0.008, binomial test) showing significant firing rate changes in response to whisker stimulation alone (4.4% decreasing firing rate, and 4.4% increasing firing rate).

By showing similar effects in awake and anesthetized mice, these recordings demonstrate that somatosensory modulation in the auditory thalamocortical system exists independently of brain state. However, while a similar level of suppression was induced by whisker deflection in awake and anesthetized mice for tones presented at 60 dB SPL, weaker effects were observed in awake animals for 80 dB SPL tones. It is possible that the higher sound level results in greater changes in the level of arousal or movement-related signals in the awake preparation that reduce the salience of the experimental whisker deflections.

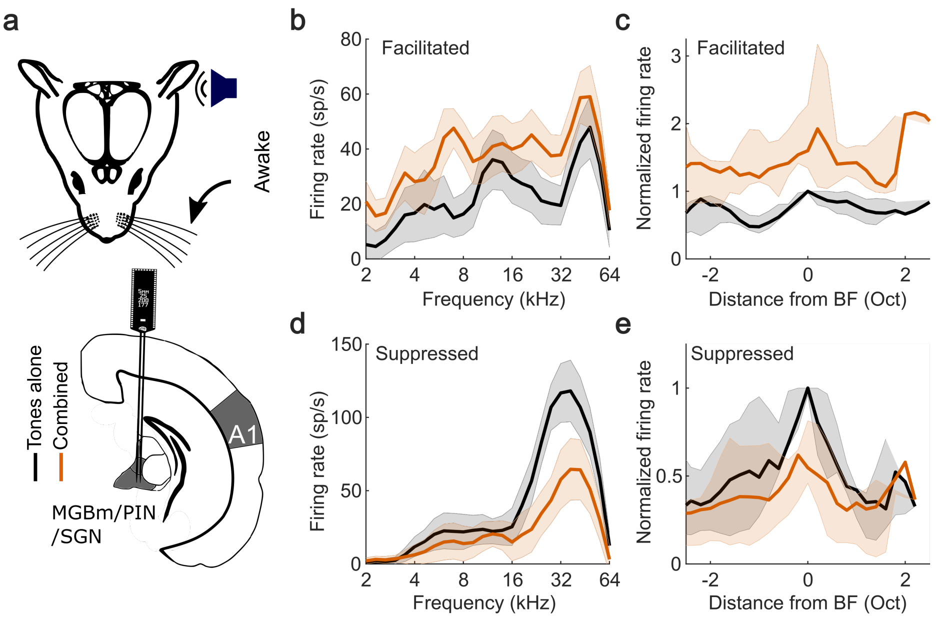


**Supplementary Fig. 6 – Related to Figure 3. Diverse somatosensory influences on neurons in MGBm/PIN and SGN of awake, passively listening mice. a** Schematic of recording setup**. b** Example frequency response profile for tones with (orange) and without (black) concurrent whisker stimulation for a unit showing crossmodal facilitation ($P < 0.05$, *t*-test). **c** Summary frequency response profiles of units with significantly facilitated BF responses (7/52 units). **d,e** Same as **b**,**c** for units with significantly suppressed BF responses (5/52). $n_{facilitated} = 7$, $n_{suppressed} = 5$, 3 mice. Shaded area indicates 95% confidence intervals of the mean (**b**,**d**) or nonparametric confidence intervals of the medians (**c**,**e**), respectively.

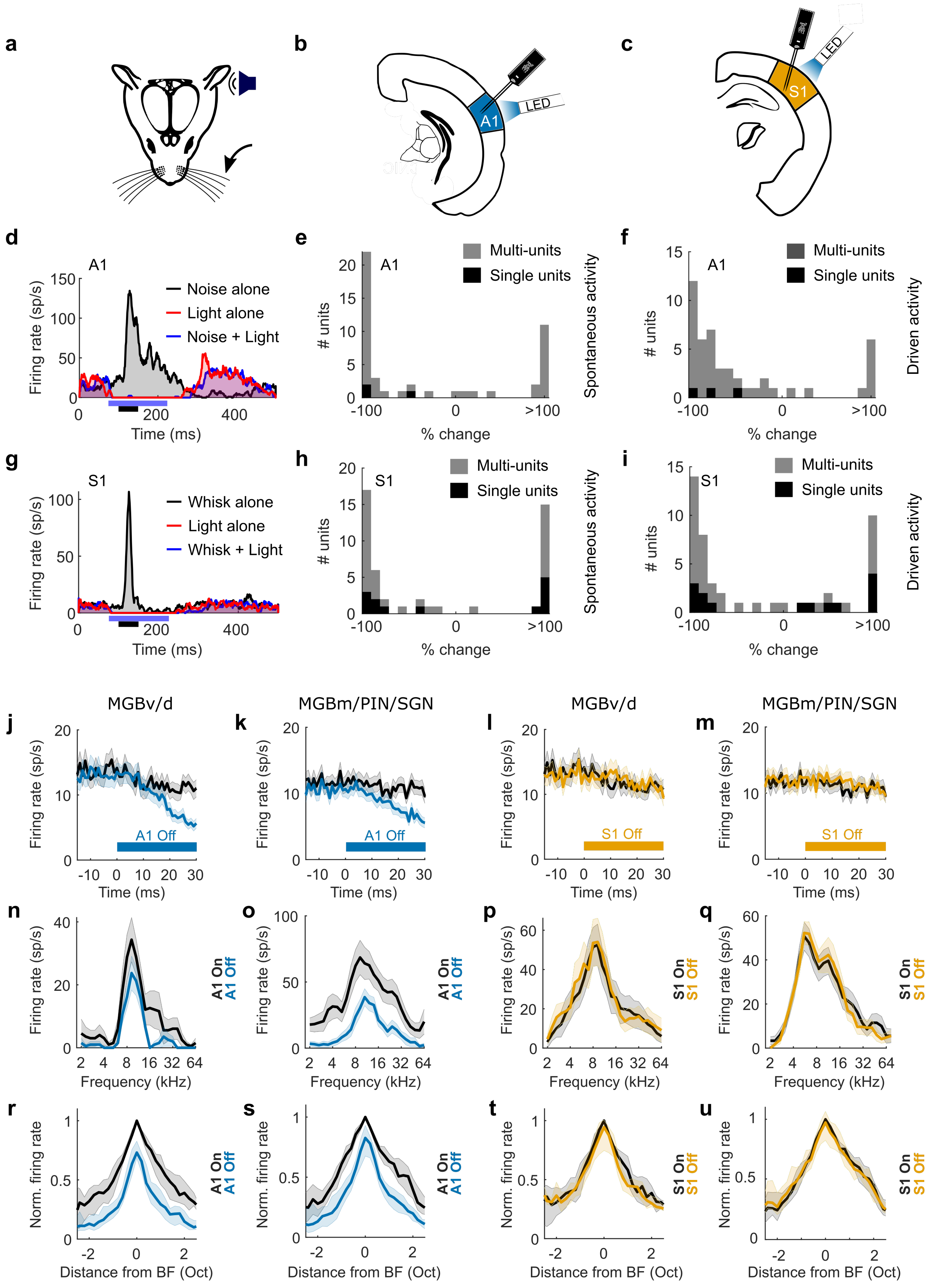


**Supplementary Figure 7 – Related to Figure 5. Effects of transient silencing of excitatory activity in auditory cortex or somatosensory cortex on cortical and thalamic activity of anesthetized mice. a** Schematic of auditory and whisker stimulation. **b** Silencing and recording of A1 activity. **c** Silencing and recording of S1 activity. **d** Example PSTHs (showing spontaneous activity and driven responses to 80 dB SPL broadband noise) from a unit in A1 during optogenetic silencing of auditory cortical activity using blue light stimulation in VGAT-ChR2-YFP mice. **e**,**f** Summary of percentage change in spontaneous activity (**e**) and noise-driven responses (**f**) in A1 following transient optogenetic silencing of excitatory cortical activity using optogenetic stimulation (ChR2) of VGAT+ cells in A1 in VGAT-ChR2-YFP mice (*n* = 47, 1 mouse). **g** Example PSTHs (showing spontaneous activity and driven responses to whisker deflection) from a unit in S1 during optogenetic silencing of somatosensory cortical activity using blue light stimulation in VGAT-ChR2-YFP mice. **h**,**i** Summary of percentage change in spontaneous activity (**h**) and whisker-driven responses (**i**) in S1 following transient optogenetic silencing of excitatory activity using optogenetic stimulation (ChR2) of VGAT+ cells in S1 in VGAT-ChR2-YFP mice (*n* = 48, 1 mouse). **j** Mean effect, across units, of A1 silencing on spontaneous activity in MGBv/MGBd. **k** Mean effect, across units, of A1 silencing on spontaneous activity in MGBm/PIN/SGN. **l** Mean effect, across units, of S1 silencing on spontaneous activity in MGBv/MGBd. **m** Mean effect, across units, of S1 silencing on spontaneous activity in MGBm/PIN/SGN. **j-m** Shaded error bars denote 95% confidence intervals around the means. Black: No light stimulation, Blue: Light stimulation starting at time 0 ms. **n** Frequency response profile of an example unit in the MGBv with (blue) or without (black) A1 silencing. **o** Frequency response profile of an example unit in the MGBm/PIN/SGN with (blue) or without (black) A1 silencing. **p** Frequency response profile of an example unit in the MGBv with (yellow) or without (black) S1 silencing. **q** Frequency response profile of an example unit in the MGBm/PIN/SGN with (yellow) or without (black) S1 silencing. **n-q** Shaded error bars denote 95% confidence intervals around the means. **r** Normalized frequency response profiles of MGBv/MGBd units with (black) or without (blue) auditory cortical silencing. **s** Normalized frequency response profiles of MGBm/PIN/SGN units with (black) or without (blue) auditory cortical silencing. **t** Normalized frequency response profiles of MGBv/MGBd units with (black) or without (yellow) S1 silencing. **u** Normalized frequency response profiles of MGBm/PIN/SGN units with (black) or without (yellow) S1 silencing. **r-u** Shaded error bars denote 95% non-parametric confidence intervals around the medians. $n_{MGBv/d}$ = 59, 3 mice; $n_{MGBm/PIN/SGN}$ = 84, 3 mice.

The spontaneous activity of noise-responsive (P < 0.001, *t*-test, *n* = 47, 1 mouse) units in A1 was strongly modulated by optogenetic activation (blue light stimulation of ChR2) of cortical VGAT+ cells in VGAT-ChR2-YFP mice. A bimodal distribution of modulation was observed. 53% of A1 units that were responsive to broadband noise at 80 dB SPL had their spontaneous activity significantly suppressed (P < 0.05, *t*-test) during blue light stimulation over A1, whereas 32% were driven by photostimulation. When removing units likely dominated by VGAT+ cell activity (i.e. units significantly driven by blue light), A1 spontaneous activity was strongly suppressed (median change = -97.9%, where -100% would indicate no spikes detected during light stimulation). Furthermore, 75% of A1 units that were responsive to broadband noise at 80 dB SPL had their responses significantly suppressed (P < 0.05, *t*-test) and 17% had their responses significantly facilitated (P < 0.05, *t*-test) during light stimulation over A1. When removing units likely dominated by VGAT+ cell activity (i.e. units with significantly facilitated responses during light stimulation), A1 responses to broadband noise (80 dB SPL) were strongly suppressed (median change = -81.5%).

Spontaneous activity of S1 units responsive to whisker deflection (P < 0.001, *t*-test, *n* = 48, 1 mouse) was strongly modulated by optogenetic activation (blue light stimulation of ChR2) of cortical VGAT+ cells in VGAT-ChR2-YFP mice. As expected, a bimodal distribution of modulation was again observed. 64.6% of single and multi-units in S1 that were responsive to whisker deflection had their spontaneous activity significantly suppressed (P < 0.05, *t*-test) during blue light stimulation over S1, while 27.6% of units were significantly driven (P < 0.05, *t*-test) by light stimulation. When removing units likely dominated by VGAT+ cell activity (i.e. units significantly driven by light), S1 spontaneous activity was strongly suppressed (median change = -92.6%). 60.4% of S1 units that were responsive to whisker deflection had their responses significantly suppressed (P < 0.05, *t*-test), while 33.3% of units had their responses significantly facilitated (P < 0.05, *t*-test) during light stimulation over S1. When removing units likely dominated by VGAT+ cell activity (i.e. units with significantly facilitated responses during light stimulation), S1 responses to whisker deflection were strongly suppressed (median change = -94.3%).

Following light stimulation of ChR2 in VGAT+ cells in A1, which has an overall suppressive effect on A1, spontaneous activity recorded in the auditory thalamus decreased in both the MGBv/MGBd (median change = -50%, P < 0.001) and the medial sector of auditory thalamus (SGN/MGBm/PIN, median change = -50%, P < 0.001). Spontaneous activity during the light-on condition started to diverge significantly (P < 0.05, t-test) from control 19 ms after light onset in the MGBv/MGBd, and 17 ms after light onset in the medial sector of auditory thalamus. Silencing A1 also decreased BF responses in both MGBv/MGBd (median change = -22.9%, P < 0.001) and in the medial sector of auditory thalamus (median change = -27%, P < 0.001). In accordance with the reduction in spontaneous activity, the smaller responses to tones appeared to reflect subtractive scaling of the frequency response profile.

In contrast to the effects of suppressing activity in A1, silencing S1 using the same optogenetic approach did not affect either spontaneous activity (P > 0.05) or BF responses (P > 0.05) in the lateral or medial sectors of the auditory thalamus. This demonstrates the specificity of these cortical manipulations, with effects of S1 silencing on the auditory thalamus observed only when somatosensory inputs from the whiskers are combined with sound.

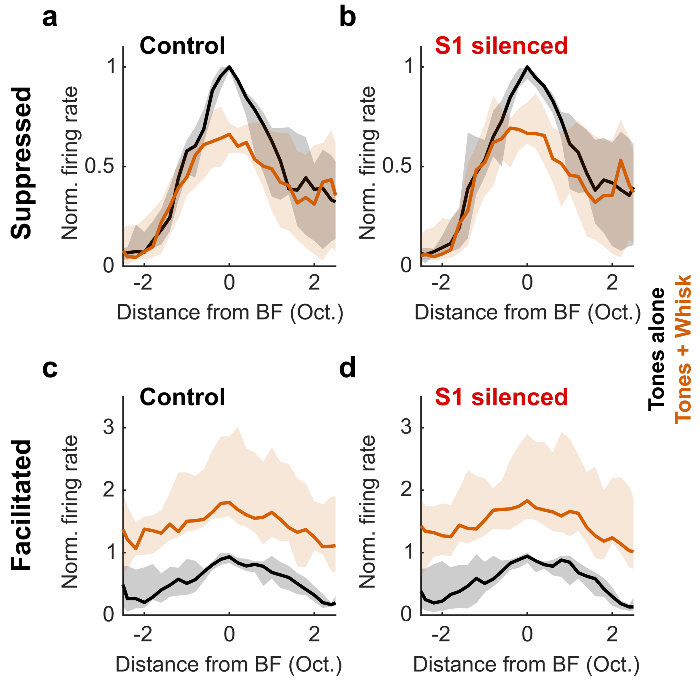


**Supplementary Figure 8 – Related to Figure 5. Silencing somatosensory cortex neither reduced whisker-driven suppression in the medial sector of auditory thalamus nor affected facilitation in anesthetized mice. a** Normalized median frequency response profiles of auditory units in MGBm/PIN and SGN (black) that were significantly suppressed ($P < 0.05$, *t*-test) by concurrent (orange) whisker deflection under control conditions. **b** As in **a** but during optogenetic silencing of S1 ($P = 0.07$, $n = 11/84$). **c** Normalized median frequency response profiles of units in MGBm/PIN and SGN (black) that were significantly facilitated ($P < 0.05$, *t*-test) by concurrent (orange) whisker deflection under control conditions. **d** As in **c** but during optogenetic silencing of S1 ($P > 0.05$, $n = 10/84$). Shaded error bars denote 95% nonparametric confidence intervals around the medians

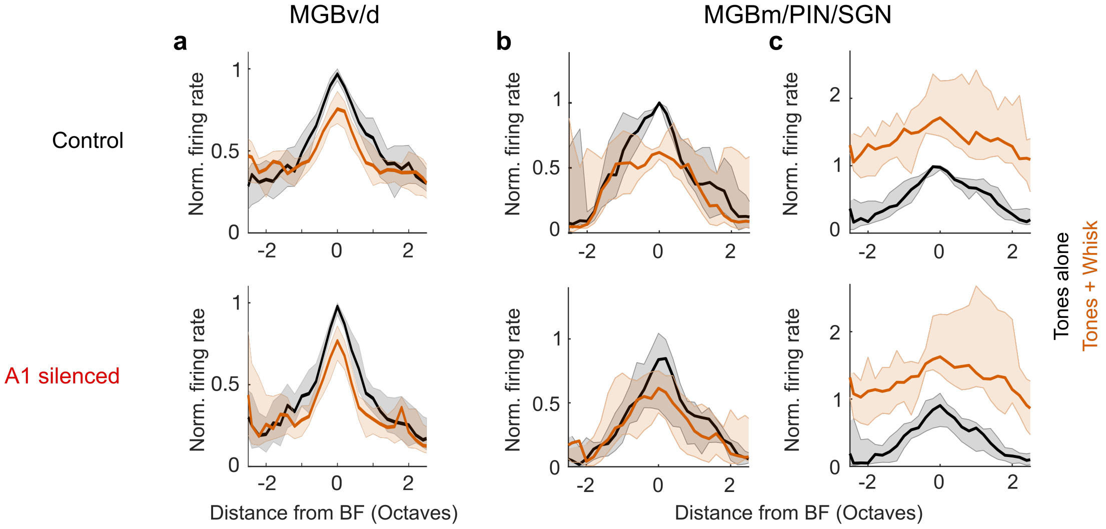


**Supplementary Figure 9 – Related to Figure 5. Somatosensory modulation of the auditory thalamus is independent of the auditory cortex in anesthetized mice. a** Normalized median frequency response profiles for all units recorded in MGBv/MGBd measured with (orange) or without (black) concurrent whisker deflection in the control condition (*top*) and with A1 silenced (*bottom*) ($n$ = 59, 3 mice). **b**,**c** Units in the medial sector of the auditory thalamus (MGBm/PIN/SGN) could be either suppressed or facilitated by somatosensory inputs. After identifying BF responses that were significantly modulated by concurrent whisker input ($P < 0.05$, *t*-test; 21/84 units, 3 mice), they were split into the two categories, suppressed or facilitated. **b** Normalized median frequency response profiles across units recorded in MGBm/PIN/SGN that showed significantly suppressed ($P < 0.05$, *t*-test) BF responses measured with (orange) or without (black) concurrent whisker deflection in the control condition (*top*) and with A1 silenced (*bottom*). **c** Normalized median frequency response profiles across units recorded in MGBm/PIN/SGN that showed significantly facilitated ($P < 0.05$, *t*-test) BF responses measured with (orange) or without whisker deflection (black) in the control condition (*top*) and with A1 silenced (*bottom*). Silencing A1 did not affect somatosensory suppression of auditory responses in MGBv/MGBd ($P > 0.05$, Wilcoxon signed-rank test), nor did it affect either the suppression or facilitation induced by whisker deflection in MGBm/PIN/SGN ($P > 0.05$, Wilcoxon signed-rank test).

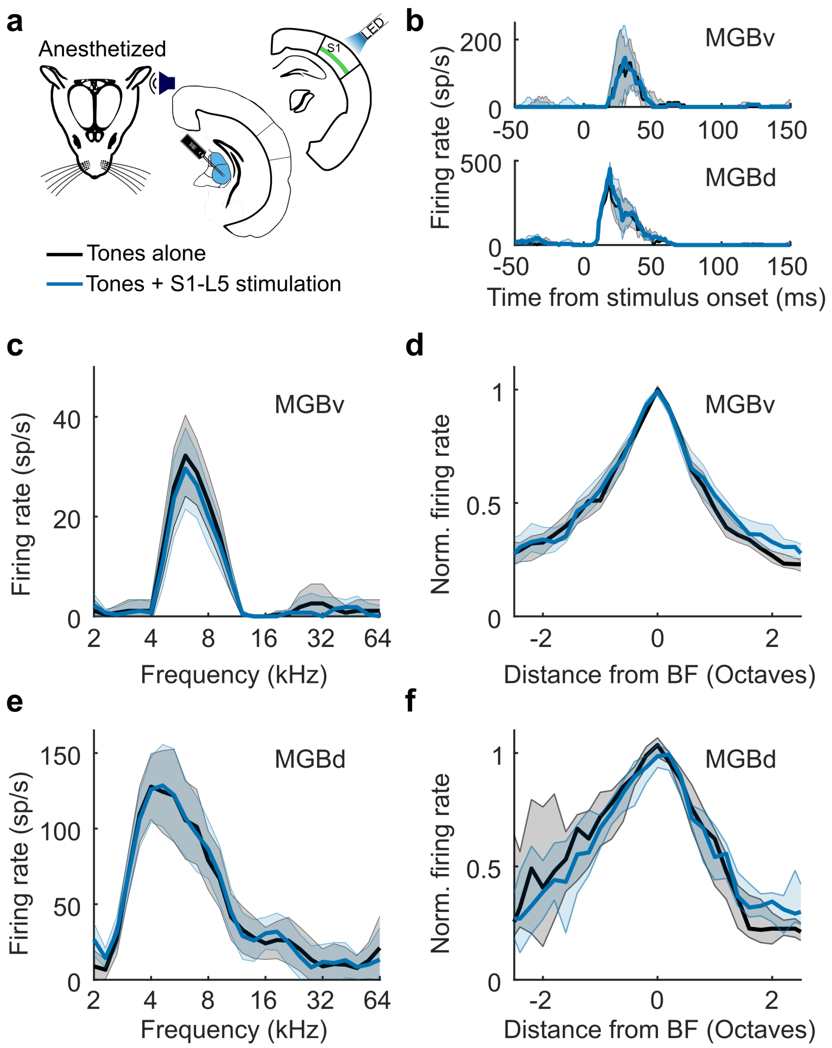


**Supplementary Figure 10 – Related to Figure 6. S1 layer 5 stimulation has no effect on auditory responses in MGBv and MGBd in anesthetized mice. a** Schematic of stimulation and recording setup. **b** Example PSTHs of BF responses from units in MGBv (*top*) and MGBd (*bottom*). **c** Frequency response profiles from the same MGBv unit. Shaded error bars denote 95% confidence intervals around the means. **d** Normalized median frequency response profiles across units in MGBv. Shaded error bars denote 95% nonparametric confidence intervals around the medians. **e**,**f** Same as **c**,**d**, but for example and pooled MGBd units. $n_{MGBv}$ = 185, 6 mice; $n_{MGBd}$ = 23, 4 mice.

Figure 6 shows that S1-layer 5 (RBP4+) cells can drive neural activity and facilitate auditory responses in the medial sector of the auditory thalamus. However, S1-L5 (RBP4+ cells) activation had no effect on the responses of units recorded in MGBv and MGBd to pure tones presented at 80 dB SPL ($P > 0.05$, Wilcoxon signed rank test).

When we examined the projection patterns from S1-bfd RBP4+ cells vs pan-neuronal S1-bfd (see the Allen Mouse Brain Atlas Experiment IDs: 127866392, 112882565, 100141473 with S1-bfd pan-neuronal labelling and 272735030, 64825323, 647806688 with S1-bfd RBP4+ labelling), we found that RBP4+ cells in S1-bfd have a very limited projection to the IC, mostly confined to its rostral sector. This is in contrast to a denser projection traversing the lateral shell of IC that is visible when labeling S1-bfd pan-neuronally. The presence of a weak input to the IC from RBP4+ neurons in layer 5 of S1-bfd likely explains why MGB responses were not suppressed by optogenetic activation of these neurons.

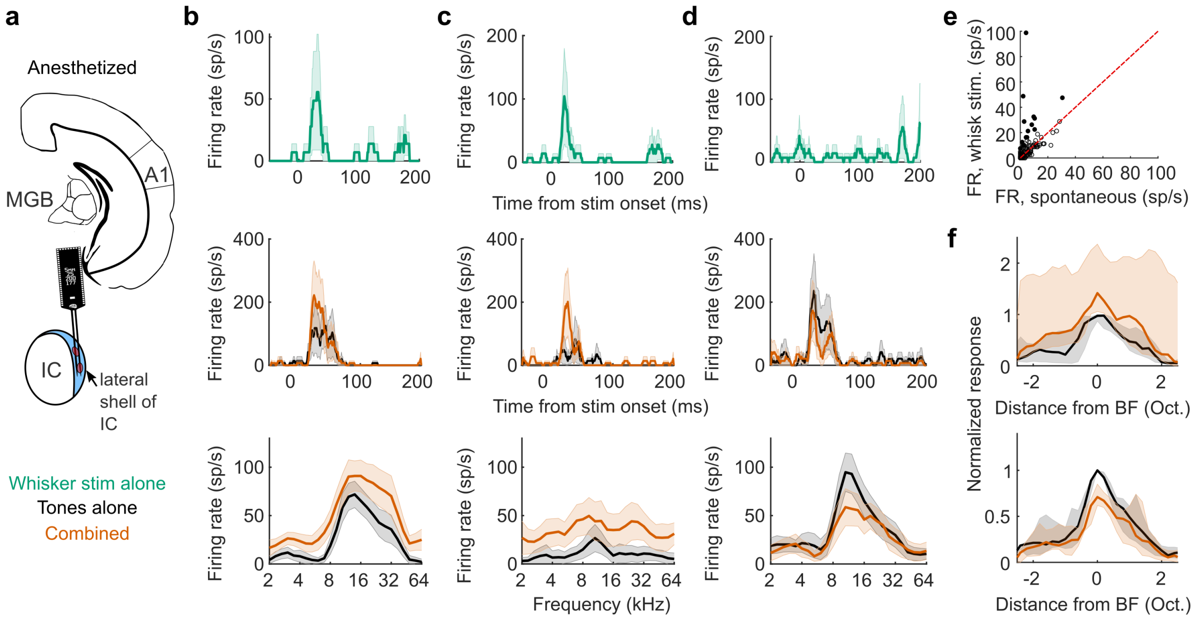


**Supplementary Figure 11 – Related to Figure 7. Neurons in the lateral shell of the IC can be driven by whisker stimulation and show somatosensory modulation of auditory responses. a** Schematic of recording setup**. b** Example unit showing response to whisker stimulation alone (*top*) and a facilitated auditory response when paired with whisker stimulation, as shown by the PSTH of the BF response (*middle*) and the frequency response profile at 80 dB SPL (*bottom*) of this neuron. **c-d** Two additional examples from recordings in the lateral shell of IC – another driven/facilitated unit (**c**) and a unit suppressed by somatosensory inputs (**d**). **e** Summary of responses to whisker stimulation alone in the lateral shell of IC. Filled circles denote significantly driven units. **f** Summary frequency response profiles of units with significantly facilitated or suppressed BF responses. $n_{\text{facilitated}}$ = 6/94, $n_{\text{suppressed}}$ = 9/94, 2 mice. Shaded area indicates 95% confidence intervals of the mean (**b,c**) or nonparametric confidence intervals of the medians (**f**).

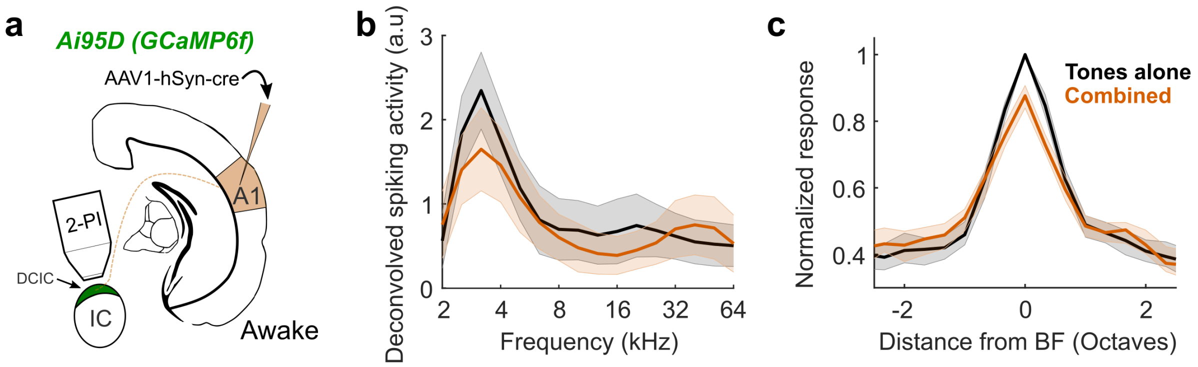


**Supplementary Figure 12 – Related to Figure 7. Dorsal cortex of the IC (DCIC) neurons receiving descending auditory cortical inputs are suppressed by whisker stimulation.** **a** Schematic of experimental setup in which A1-recipient neurons in the optically accessible DCIC were labelled by cortical injection of AAV1-hSyn-cre virus**. b** Frequency response profiles obtained by 2-photon imaging in an awake mouse of an example A1-recipient IC neuron with (orange) or without (black) concurrent whisker deflection. **f** Median frequency response profiles across all tuned A1-recipient neurons in the dorsal cortex of the IC with (orange) or without (black) concurrent whisker deflection (*n* = 232 cells, 2 mice). Shading illustrates the 95% confidence intervals around the mean (**b**) or nonparametric 95% confidence intervals around the median firing rate (**c**).

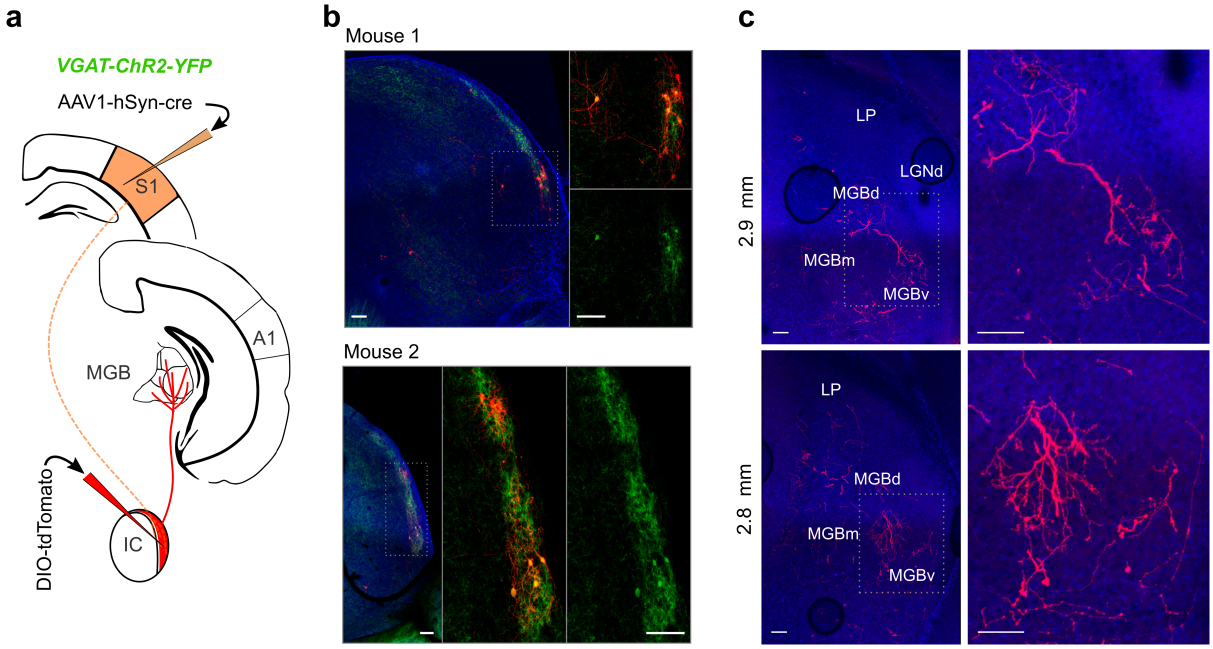


**Supplementary Figure 13 – Related to Figure 7. S1-recipient IC shell neurons are GABAergic and project to MGB. a** Schematic of experiment in which S1-recipient neurons in the shell of the IC were labelled trans-synaptically. **b** Examples of co-labelling of GABAergic neurons (*VGAT*-ChR2-eYFP) and S1-recipient neurons (tdTomato) in the lateral shell of IC in two example mice. The labelling suggests that at least a substantial proportion of S1-recipient LCIC neurons are GABAergic. The areas enclosed by the white dotted boxes in the left images are shown at high magnification on the right, with and without tdTomato labelling of S1-recipient cells. **c** Examples of axons (tdTomato) from S1-recipient IC neurons in the auditory thalamus. The areas enclosed by the white dotted boxes in the left images are shown at high magnification on the right. Blue is DAPI cell staining. Scale bars = 100 μm.

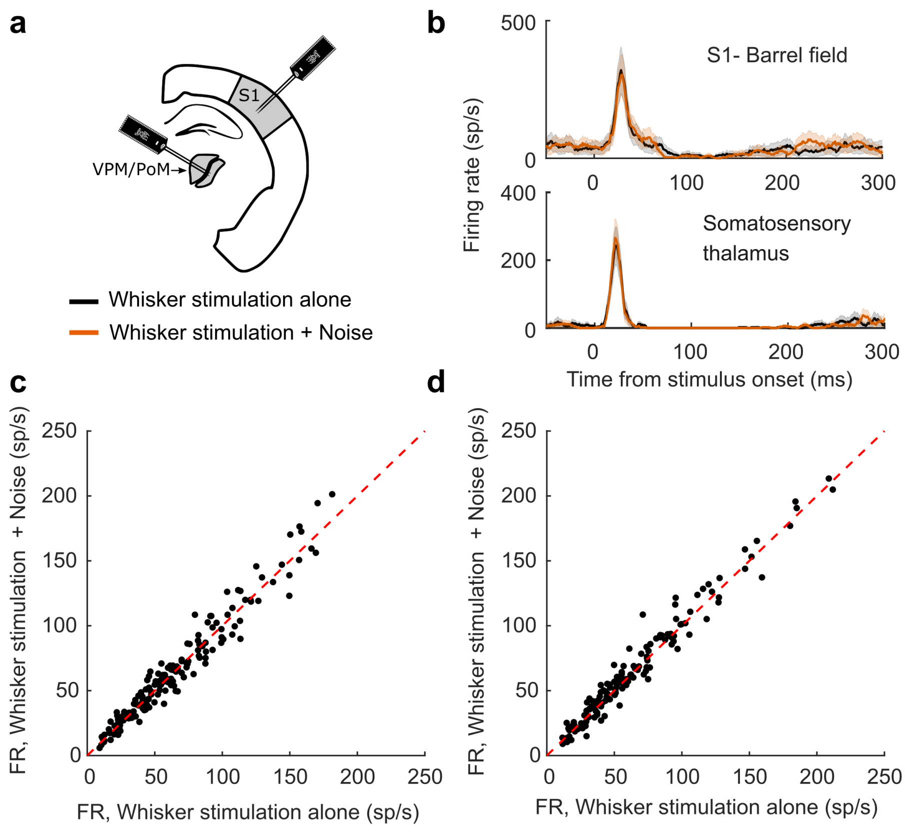


**Supplementary Figure 14 – Related to Supplementary Figure 4. Minimal auditory modulation of somatosensory responses in somatosensory thalamus and S1 of anesthetized mice. a** Schematic of recording setup. **b** Examples of responses to whisker stimulation of units recorded in the barrel field of S1 (*top*) and somatosensory thalamus (*bottom*) with (orange) or without (black) concurrent noise bursts at 80 dB SPL. **c** Summary of all units recorded in the barrel field of S1 that responded to whisker deflection, showing firing rate evoked by whisker deflection alone vs whisker deflection with concurrent noise bursts. **d** As in **c**, but for all units recorded in the somatosensory thalamus.

To assess the reciprocity of the somatosensory influences on the auditory thalamocortical system, extracellular multi- and single units were recorded in the vibrissae somatosensory thalamus (ventroposterior medial nucleus (VPM) and posterior medial nucleus (PoM)) in response to 50-ms whisker stimulation, with (orange) or without (black) concurrent 50-ms broadband noise stimulation at 80 dB SPL (i.e. the same stimuli used in the recordings from the auditory system). No effect of auditory stimulation was found on responses in the barrel field of S1 (Wilcoxon signed rank tests, $P > 0.05$, $n = 170$, 4 mice). We found a slight, but significant, increase in firing rate during combined auditory-somatosensory stimulation, relative to somatosensory stimulation alone, in somatosensory thalamus (Wilcoxon signed rank tests, $P = 0.01$, $n = 154$, 5 mice). Sound stimulation alone did not induce any effects on baseline firing rate in S1 or somatosensory thalamus.
